# High-Throughput Screening Identifies Small-Molecule Inhibitors of the Tau-LRP1 Interaction

**DOI:** 10.64898/2026.06.24.733881

**Authors:** Caiqin Wang, Chen-Ting Ma, Camryn Crotty, Fu-Yue Zeng, Andrey Bobkov, Jonathan A. Covel, Erica Keane Rivera, Eduard Sergienko, Kenneth S. Kosik, Steven H. Olson, Michael R. Jackson, Jennifer N. Rauch

## Abstract

The cellular uptake and propagation of tau are central features of tauopathies, including Alzheimer’s disease, and are mediated by the endocytic receptor low-density lipoprotein receptor-related protein 1 (LRP1). While prior studies have implicated LRP1 in tau binding and internalization, the biochemical features of this interaction and its suitability for therapeutic targeting remain incompletely defined. Here, we establish a quantitative and scalable framework to interrogate the tau-LRP1 interaction and identify small-molecule modulators. We engineered and purified the LRP1 ligand-binding domain 4 (BD4), a key region mediating tau interaction, and developed multiple orthogonal assays, including fluorescence polarization, split luciferase complementation, and time-resolved FRET, to measure LRP1-BD4 interactions with tau and a known peptide ligand. Across assay formats, we observe consistent binding affinities in the nanomolar range and demonstrate competitive displacement by tau, receptor-associated protein (RAP), and a peptide ligand, supporting overlapping binding interfaces. Leveraging these platforms, we performed small molecule high-throughput screening and identified a set of candidate inhibitors of the LRP1-BD4-tau interaction. Selected compounds reduced tau uptake in a cellular assay, phenocopying competitive inhibition by tau and a peptide ligand. Together, these studies define the LRP1-BD4-tau interaction as a biochemically tractable and druggable interface and establish an integrated discovery pipeline linking mechanistic characterization to functional cellular outcomes. This work provides a foundation for the development of therapeutic strategies targeting LRP1-mediated tau uptake.

## Introduction

The aggregation and spread of tau protein into intracellular inclusions is a defining pathological feature of Alzheimer’s disease (AD) and related tauopathies. In AD, tau pathology progresses through the brain in a stereotyped and hierarchical manner, affecting regions critical for learning and memory, and closely correlates with cognitive decline^1,2^. A substantial body of evidence now supports a model in which the intercellular transfer of tau species drives this progression, implicating tau uptake as a key step in disease propagation^3–5^. Despite this, there are currently no effective therapeutic strategies that directly target the cellular mechanisms underlying tau spread.

The endocytic receptor low-density lipoprotein receptor-related protein 1 (LRP1) has been identified as a major mediator of tau uptake^6–8^. LRP1 facilitates tau’s internalization across multiple tau conformational states, including both monomeric and larger soluble aggregate species^6^. Importantly, LRP1 knockdown reduces tau uptake in cells, reduces cellular seeding, and prevents tau spread *in vivo*^6,7^, establishing the tau-LRP1 interaction as a critical node in disease progression and a compelling target for therapeutic intervention. However, the large size and structural complexity of LRP1, a ∼600kDa receptor composed of multiple ligand-binding domains^9,10^, has limited the detailed biochemical interrogation of this interaction.

Previous work from our group demonstrated that the fourth ligand-binding domain of LRP1 (BD4) is sufficient to mediate tau uptake in cell culture, identifying this region as a minimal functional unit for tau recognition^6^. BD4 is composed of 11 serial complement-type repeats that engage ligands via electrostatic interactions in a calcium-dependent fashion^11^. Prior work has shown that the receptor-associated protein (RAP) can function as a pan-LRP1 antagonist that competitively blocks all known ligands^12,13^ and engineered RAP variants have demonstrated LRP1 inhibition both *in vitro* and *in vivo*^14^. These properties suggest that the tau-LRP1 interaction may be amenable to pharmacological modulation. However, efforts to directly target this interface have been hindered by the lack of robust recombinant systems for LRP1 and the absence of quantitative, screening-compatible assays to measure its interaction with tau.

From a chemical biology perspective, the development of tractable assay platforms is essential for enabling the discovery of small-molecule modulators of protein-protein interactions. In the case of LRP1, ligand binding is mediated by multivalent and electrostatic interactions^15–17^, which complicates assay design and increases susceptibility to non-specific inhibition in screening campaigns. These challenges necessitate the use of orthogonal assay formats that can quantitatively measure binding while enabling rigorous assessment of compound mechanism and specificity.

Here, we report the recombinant production and biochemical characterization of LRP1 binding domain 4 (LRP1-BD4) and the development of a suite of orthogonal assays to quantify its interactions with tau and other established LRP1 ligands. Using these platforms, including fluorescence polarization, split luciferase complementation, and time-resolved FRET, we establish sensitive and scalable methods to interrogate the tau-LRP1 interaction. We perform two high-throughput screening campaigns to identify small-molecule inhibitors and demonstrate that selected compounds reduce tau uptake in cells. Together, this work establishes a biochemical and screening framework for targeting LRP1-mediated tau uptake and provides a foundation for the development of chemical probes to study and modulate tau propagation.

## Results and Discussion

### Recombinant production and biochemical characterization of LRP1-BD4

To enable biochemical interrogation and eventual small molecule screening of the tau-LRP1 interaction, we generated a recombinant mammalian expression construct encoding the LRP1 ligand binding domain 4 (BD4; residues 3293-3783), which was previously implicated in tau binding^6^. The LRP1-BD4 construct was designed with an N-terminal Maltose Binding Protein (MBP) tag to enhance solubility, followed by tandem TEV and thrombin cleavage sites, a Flag epitope tag, the BD4 domain, and a C-terminal His tag to facilitate purification and detection (**Figure 1A**). Expression of LRP1-BD4 was achieved with co-expression of the LRP1 molecular chaperone RAP in mammalian Expi293F cells, which yielded robust levels of secreted protein. To purify LRP1-BD4 we performed affinity chromatography in the presence of 100mM EDTA to remove bound RAP. After purification, protein was dialyzed back into Ca^+2^ containing buffer and efficient removal of the MBP tag was achieved via thrombin cleavage. Efficient cleavage was confirmed by SDS-PAGE, which demonstrated the expected shift in molecular weight and high purity of the cleaved BD4 domain (**Supplementary Figure 1A**).

**Figure 1.**
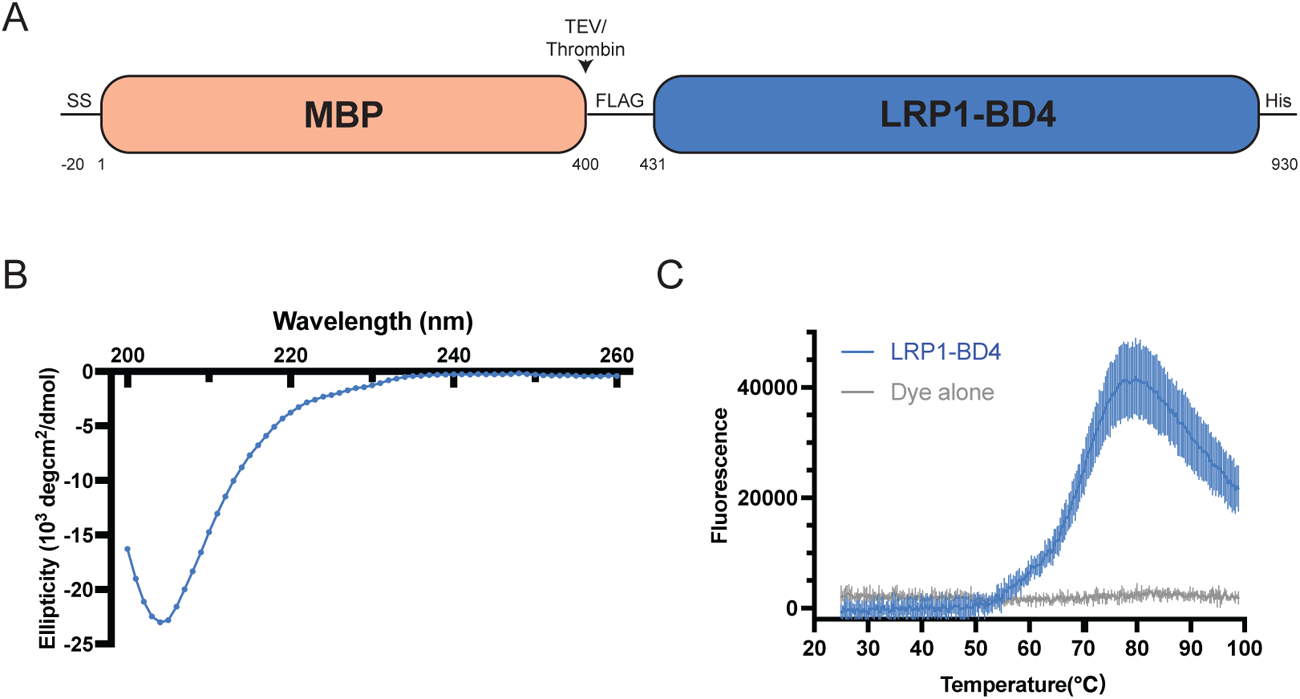
Recombinant production and biochemical characterization of LRP1-BD4. (A) Schematic of the LRP1-BD4 expression construct. Construct contains an N-terminal MBP tag, followed by TEV/thrombin cleavage sites, Flag tag, BD4 domain (residues 3293-3783), and a C- terminal His tag. SS = signal sequence for proper ER targeting. (B) Circular dichroism spectra of purified LRP1-BD4 from 200-260nm. (C) Representative DSF thermal melt curve of LRP1-BD4 (blue) vs. dye alone (grey) (mean±SD, technical triplicates). T_m_ = 68.1 ± 0.5°C, average of n=2 independent experiments.

To assess structural integrity of the produced LRP1-BD4, we performed circular dichroism (CD). CD analysis revealed low alpha-helical and beta-sheet content (**Figure 1B**), consistent with the structural properties of complement-type repeats (CR) that make-up the BD4 domain^18^ as well as structural predictions from AlphaFold3 (**Supplementary Figure 1B**). Thermal stability was evaluated using differential scanning fluorimetry (DSF), which showed a melting temperature of 68.1 +/- 0.5°C (**Figure 1C**), indicating that the isolated LRP1-BD4 is quite thermally stable. Again, this is consistent with known features of the BD4 domain, which is stabilized by both a conserved network of 33 intradomain disulfide bonds and calcium coordination at each repeat^19^.

### Development of a fluorescence polarization assay to quantify LRP1-BD4 ligand interactions

To determine whether the produced LRP1-BD4 protein was functional, we established a fluorescence polarization (FP) assay to quantitatively measure ligand binding to LRP1-BD4. L57, an artificial peptide discovered via phage display to bind specifically to the fourth ligand binding domain of LRP1 (BD4)^20^, was purchased conjugated to FITC to serve as a fluorescent tracer. Incubation of FITC-L57 (20nM) with increasing concentrations of LRP1-BD4 resulted in a concentration-dependent increase in polarization, enabling determination of binding affinity. Analysis of the binding curve yielded an apparent dissociation constant K_d_ of 42nM (95% CI: 34- 51nM, n=5) (**Figure 2A**), consistent with previous ELISA measurements of L57 and LRP1 binding^20^.

**Figure 2.**
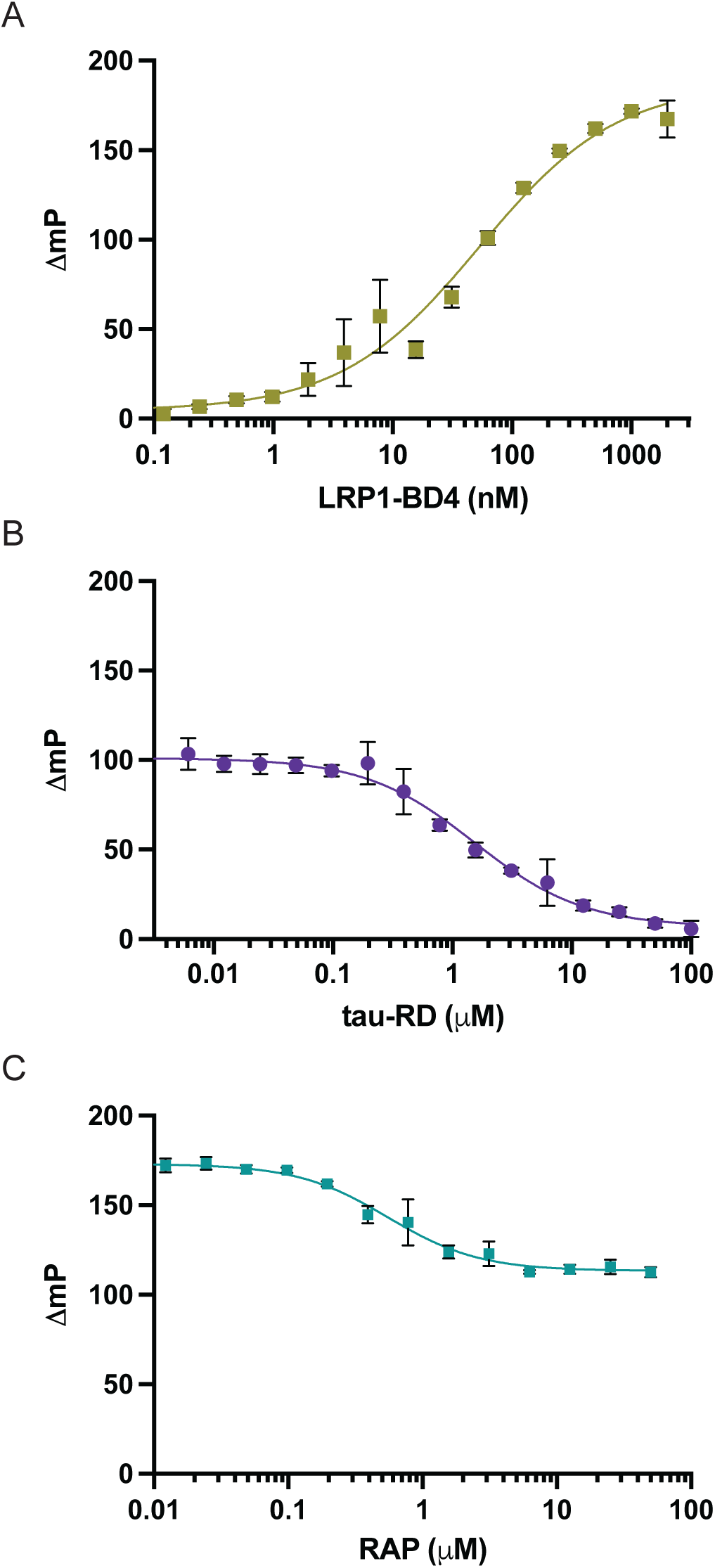
Fluorescence polarization assay quantifies LRP1-BD4 ligand binding. (A) Direct binding of FITC-L57 to LRP1-BD4. Increasing concentrations of LRP1-BD4 were incubated with 20nM FITC-L57 and polarization was measured, representative experiment is shown (mean±SD, technical triplicates), K_d_ = 42nM (95% CI: 34-51nM, n=5 independent experiments). (B) LRP1-BD4 (1μM) / FITC-L57 (20nM) competition with tau-RD, representative experiment is shown (mean±SD, technical triplicates), IC_50_ = 1.6μM (95% CI:1.2-2.1μM, n=3 independent experiments). (C) LRP1-BD4 (1μM) / FITC-L57 (20nM) competition with RAP, representative experiment is shown (mean±SD, technical triplicates), IC_50_ = 0.8μM (95% CI:0.4-1.4μM, n=4 independent experiments).

To evaluate assay specificity, we performed competition experiments using unlabeled L57, tau microtubule-repeat domain construct (“tau-RD”; residues 244-372) which was previously shown to utilize LRP1 for endocytosis^6^, and RAP. Competition of 100nM LRP1-BD4/20nM L57-FITC with unlabeled L57 peptide yielded an IC_50_ of 156nM (95% CI: 104-234nM, n=3) and a calculated K_i_ of 134nM (**Supplementary Figure 2A**). The K_i_ is approximately 3-fold higher than the K_d_ (42nM) measured by direct tracer titration and may reflect the contribution of the FITC label to the measured binding. Based on our knowledge that LRP1 engages lysine residues on ligands for binding, we purchased an L57 peptide with all four lysine residues converted to alanine (“L57A”). L57A was not able to displace L57-FITC (**Supplementary Figure 2B**), further confirming the specificity of our assay design.

To maximize signal in our competition assays, we held LRP1-BD4 at 1μM and L57-FITC at 20nM, and performed competition experiments with both tau-RD and RAP. Tau competed with an IC_50_ of 1.6μM (95% CI:1.2-2.1μM, n=3; **Figure 2B**), whereas RAP competed with an IC_50_ of 0.8μM (95% CI: 0.4-1.4μM, n=4; **Figure 2C**). Under these conditions, the IC_50_ values for both tau and RAP exceeded the previously reported affinities for these protein-protein interactions (tau-LRP1; 60 ± 8nM^21^, RAP-LRP1; 9 ± 5nM^22^), which is consistent with tight-binding conditions and [LRP1-BD4] >> K_i_. Interestingly, while tau-RD was capable of reducing the polarization signal to baseline at saturating concentrations, RAP only partially displaced the FITC-L57 tracer. We confirmed that RAP itself does not have measurable binding to the FITC-L57 tracer (**Supplementary Figure 2C**), therefore these results indicate that RAP reduces but does not fully abolish L57 binding to LRP1-BD4, consistent with overlapping but non-identical binding sites.

### A split luciferase complementation assay provides orthogonal measurement of LRP1- BD4-tau binding

To establish an orthogonal assay, and to allow for measurement of the interaction between LRP1-BD4 and tau, we developed a split luciferase complementation system based on NanoLuc technology^23^. For this assay, we engineered an LRP1-BD4 construct containing the LgBiT tag inserted between the MBP and BD4 domains. Complementary full-length (2N4R isoform) tau constructs were generated to include the 11-amino acid SmBiT tag (N-term; “SmBiT-tau” and C-term; “tau-SmBiT”). The intrinsic affinity between LgBiT and SmBiT is weak (reported K_d_ = 190μM^23^), such that binding of LRP1-BD4 to tau brings the tags into proximity, driving luciferase reconstitution and luminescent signal generation (**Figure 3A**). Due to tau’s propensity for promiscuous binding, we also created an MBP-LgBiT construct lacking the LRP1- BD4 domain to assess any non-specific interactions due to either the MBP or LgBiT tag.

**Figure 3.**
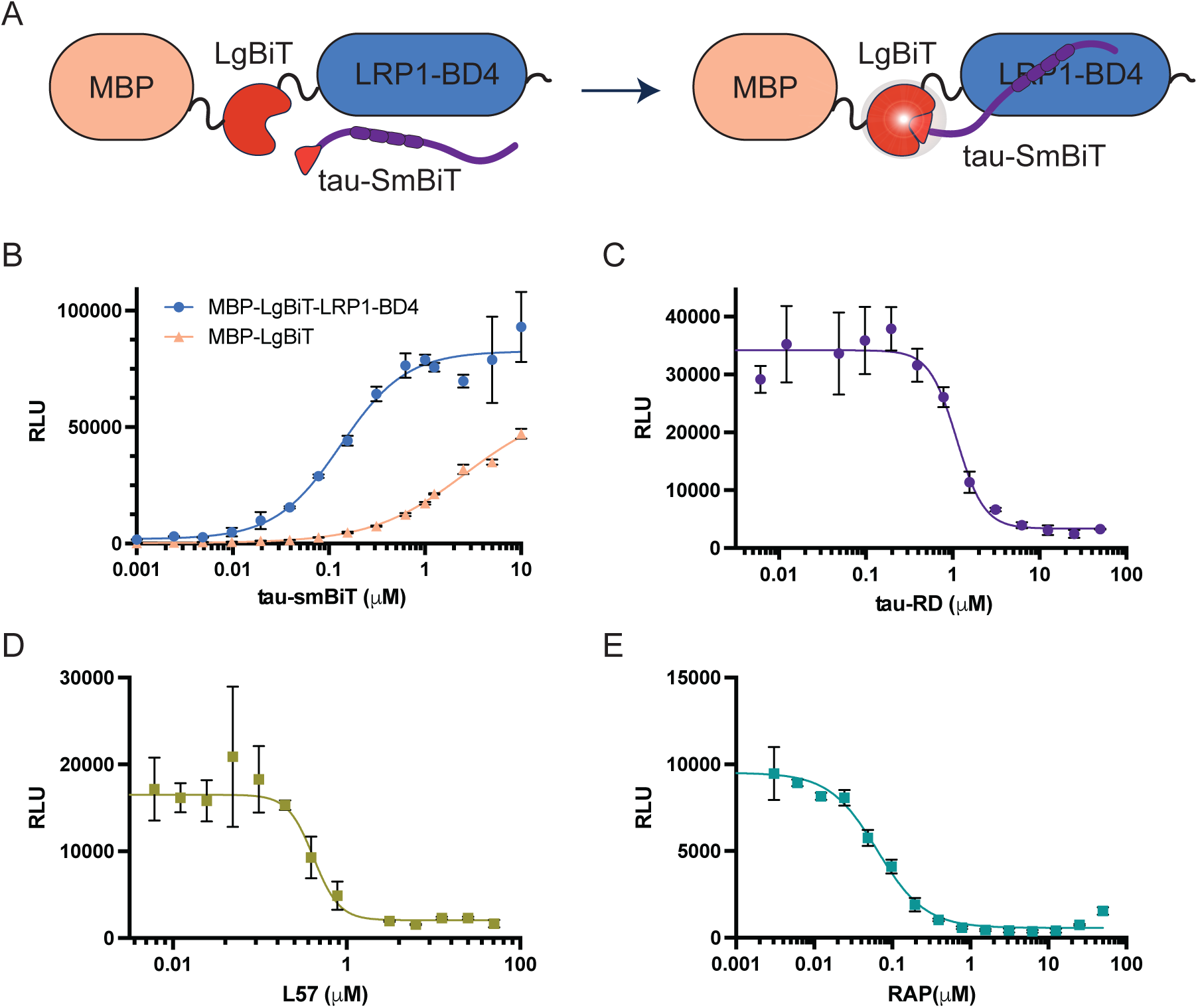
Split luciferase complementation assay permits measurement of LRP1-BD4-tau binding. (A) Schematic of the NanoBiT assay. LgBiT is inserted between MBP and BD4 domains of LRP1-BD4; SmBiT is fused to tau (N- or C-terminal). Binding of LRP1-BD4 to tau brings LgBiT and SmBiT into proximity, reconstituting luciferase activity. (B) Titration of tau- SmBiT with 10nM MBP-LgBiT-LRP1-BD4 or MBP-LgBiT, representative experiment is shown (mean±SD, technical triplicates), EC_50_ = 130nM (95% CI: 120-150nM, n=3 independent experiments). (C) Competition with untagged tau-RD. MBP-LgBiT-LRP1-BD4 (10nM) and tau- SmBiT (200nM) held constant; representative experiment is shown (mean±SD, technical triplicates). IC_50_ = 1.2μM (95% CI: 0.7-2μM, n=3 independent experiments). (D) Competition with L57. MBP-LgBiT-LRP1-BD4 (10nM) and tau-SmBiT (200nM) held constant; representative experiment is shown (mean±SD, technical triplicates). IC_50_ = 0.3μM (95% CI: 0.2-0.6μM, n=4 independent experiments). (E) Competition with RAP. MBP-LgBiT-LRP1-BD4 (10nM) and tau- SmBiT (200nM) held constant; representative experiment is shown (mean±SD), technical triplicates). IC_50_ = 56nM (95% CI: 36-87nM, n=3 independent experiments).

Consistent with specific complex formation, titration experiments revealed a concentration-dependent increase in luminescence for tau-SmBiT with MBP-LgBiT-LRP1-BD4 (**Figure 3B**) with an EC_50_ of 130nM (95% CI: 120-150nM, n=3). The control MBP-LgBiT protein showed minimal signal at tau concentrations below 1μM, confirming specificity of the tau-LRP1 interaction. The measured EC_50_ for our tau-SmBiT with MBP-LgBiT-LRP1-BD4 is approximately 2-fold higher than the previously reported K_d_ for 2N4R tau with LRP1 determined via surface plasmon resonance (60 ± 8nM^21^). The difference in affinities could be driven by either the requirement for productive LgBiT-smBiT complementation in addition to binding, such that not all binding events produce luminescence or could be a consequence of the differences in LRP1 constructs utilized across studies. Both orientations of the SmBiT tag (N-term & C-term) were capable of complementation (**Supplementary Figure 3A**).

To evaluate assay specificity, we performed competition experiments using untagged tau and L57. Holding MBP-LgBiT-LRP1-BD4 and tau-SmBiT constant (10nM and 200nM, respectively), untagged tau-RD yielded an IC_50_ of 1.2μM (95% CI: 0.7-2μM, n=3) (**Figure 3C**). L57 also displaced tau-SmBiT in a dose-dependent manner with an IC_50_ of 0.3μM (95% CI: 0.2-0.6μM, n=4) (**Figure 3D**). These results were again consistent with L57 and tau being competitive LRP1-BD4 binding partners. We also measured RAP competition in this platform and found that RAP was capable of displacing tau-SmBiT from MBP-LgBiT-LRP1-BD4 with an IC_50_ of 56nM (95% CI: 36-87nM, n=3) (**Figure 3E**). Similar competition for tau-RD, L57, and RAP was also achieved using the N-term tagged construct (smBiT-tau; **Supplementary Figure 3B**).

### TR-FRET assay platforms enable high-throughput interrogation of LRP1-BD4 interactions

To further expand our assay toolkit and enable high-throughput screening (HTS), we developed time-resolved Förster resonance energy transfer (TR-FRET) assays to monitor LRP1-BD4 interactions with both tau and the L57 peptide. TR-FRET has several advantages over other protein-protein interaction assays, including a high signal-to-noise ratio, reduced background from compound autofluorescence and fewer artifact hits^24,25^, which makes TR-FRET well suited for HTS applications. In the tau-based configuration, an Eu^3+^-labeled anti-Flag antibody served as the fluorescence donor molecule recognizing LRP1-BD4, while tau labeled with Alexa Fluor- 647 (“tau-647”) served as the acceptor (**Figure 4A**). Optimal concentrations of tau-647 and LRP1-BD4 were determined by performing a matrix experiment where concentrations of each binding partner were varied and tested (**Figure 4B**). The most robust TR-FRET signal was achieved with 10-80nM LRP1-BD4 and 40-80nM tau-647. To establish specificity of signal, unlabeled tau and L57 were used as competitors holding both tau-647 and LRP1-BD4 constant at 5nM, at which concentration assay sensitivity was maintained by keeping both protein and tracer below apparent EC_50_. Tau and L57 had IC_50_ values of 3.3nM (95% CI: 2.9-3.9nM, n=2) and 36nM (95% CI: 23-66nM, n=2) respectively (**Supplementary Figure 4A**), confirming specificity and suitability of the TR-FRET platform for screening applications.

**Figure 4.**
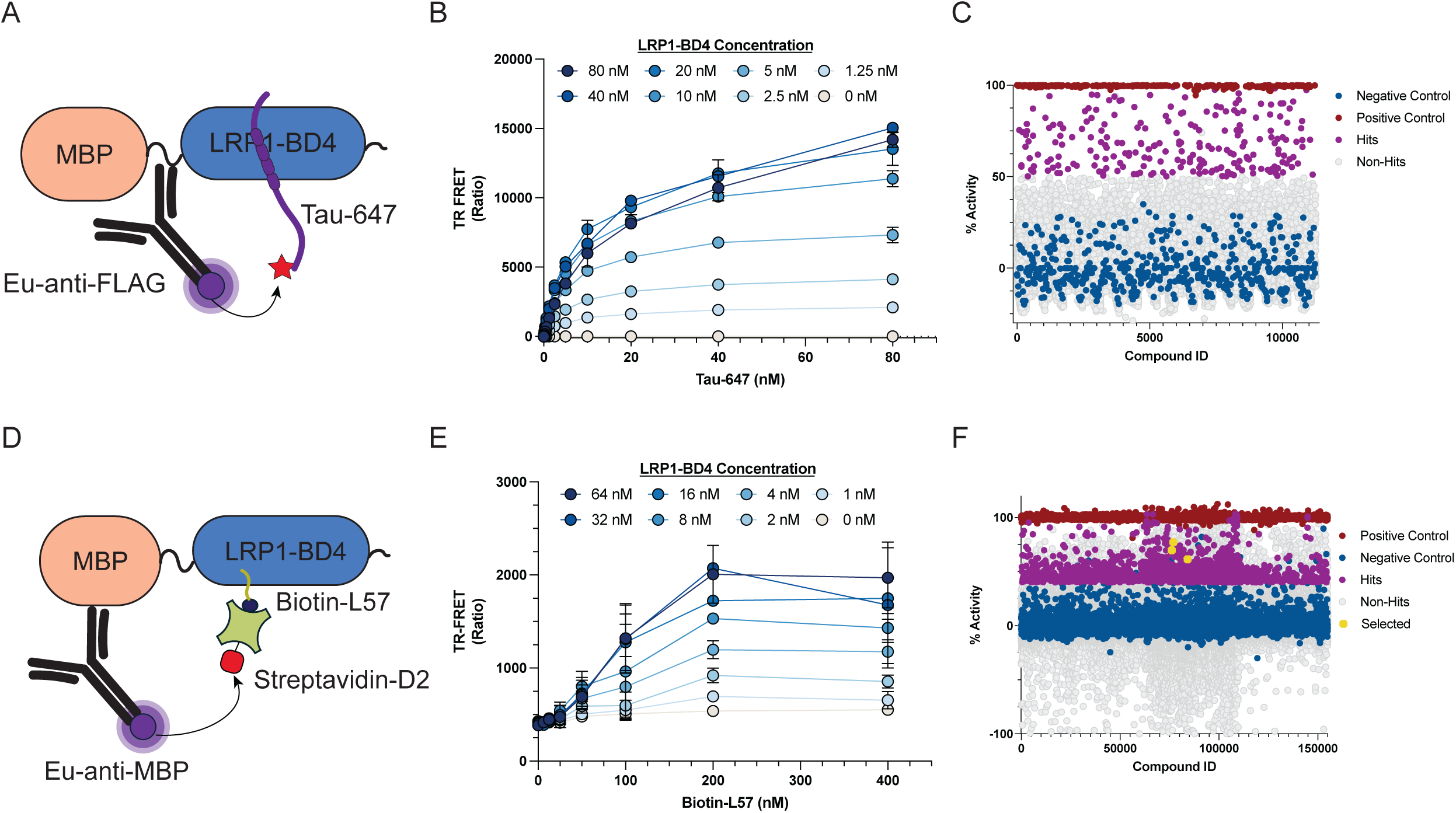
TR-FRET assay platforms facilitate high-throughput screening for molecules that disrupt LRP1-BD4 ligand interactions. (A) Schematic of the LRP1-BD4-tau TR-FRET assay. Eu^3+^-anti-Flag antibody (donor) binds LRP1-BD4; tau-647 (acceptor) binds BD4; FRET signal reports on complex formation. (B) Concentration matrix for LRP1-BD4-tau TR-FRET optimization (mean±SD, technical triplicates). (C) Scatter plot of the LRP1-BD4-tau TR-FRET primary screen of 10,240 compounds (Chembridge Library) at 25μM. X-axis is the compound ID and the y-axis is the % Activity of the compound with 100% being no LRP1-BD4 and 0% being DMSO. Negative controls are colored in blue, positive controls are in red, non-hits are shown in grey and hits are shown in purple. (D) Schematic of the LRP1-BD4-L57 TR-FRET assay. Eu^3+^- anti-MBP antibody (donor) binds LRP1-BD4; streptavidin-D2 (acceptor) detects biotinylated L57. (E) Concentration matrix for LRP1-BD4-L57 TR-FRET optimization (mean±SD, technical triplicates). (F) Scatter plot of the LRP1-BD4-L57 TR-FRET primary screen of 150,000 compounds at 25μM. X-axis is the compound ID and the y-axis is the % Activity of the compound with 100% being no Biotin-L57 and 0% being DMSO. Negative controls are colored in blue, positive controls are in red, non-hits are shown in grey, hits are shown in purple and selected compounds for follow-up work are shown as yellow stars.

Using the LRP1-BD4-tau TR-FRET assay, we screened a library of 10,240 small molecules from the Chembridge Library collection at 25μM compound concentration. While the assay exhibited strong performance metrics, the screen yielded a relatively large number of initial hits (232 compounds, 2.2% hit rate; **Figure 4C**). Although the majority of these hits re-confirmed upon cherry-picking (**Supplementary Figure 4B**), the elevated hit rate itself is a warning sign for systematic artifacts. We attribute the high hit rate to the use of tau as a binding partner in the assay. Tau, as an intrinsically disordered protein with a high propensity for promiscuous interactions, is susceptible to compound-induced aggregation or non-specific displacement which would generate reproducible but mechanistically irrelevant hits. Rather than pursuing this hit set, we developed an alternative screening strategy using the L57 peptide as a partner.

To facilitate our second HTS screening strategy, we developed an LRP1-BD4-L57 TR-FRET assay. Because we had determined that L57 and tau are competitive for binding to LRP1-BD4, we reasoned that inhibitors identified in this format would still report on disruption of the LRP1- BD4-tau interaction, while reducing enrichment of non-specific hits. In this assay system, an Eu^3+^-labeled anti-MBP antibody was used as the donor and streptavidin-D2 as the acceptor to detect interactions between LRP1-BD4 and biotinylated L57 (**Figure 4D**). Similar to the LRP1- BD4-tau TR-FRET assay, optimal concentrations of Biotin-L57 and LRP1-BD4 were determined by performing a matrix experiment where concentrations of each binding partner were varied and tested (**Figure 4E**). The most robust TR-FRET signal was achieved with 16-64nM LRP1- BD4 and 200-400nM Biotin-L57. To establish specificity of signal, unlabeled L57 and tau were used as competitors holding Biotin-L57 and LRP1-BD4 constant at 106nM and 8nM respectively. Unlabeled L57 had an IC_50_ of 106nM (95% CI: 71-167nM, n=2), unlabeled tau competed with an IC_50_ of 66nM (95% CI: 50-87nM, n=2), and L57A was non-competitive (**Supplementary Figure 4C**), validating specificity and suitability of this TR-FRET platform for HTS.

Using the LRP1-BD4-L57 platform, we screened 150,000 small molecules (25μM) in 1536-well format, holding Biotin-L57 and LRP1-BD4 constant at 200nM and 30nM respectively. The overall screen performance was excellent with S/B = 5.68 ± 0.26 and a Z-factor of 0.81 ± 0.03. An initial hit rate of 1476 compounds (0.98%) was achieved. Hits were confirmed via cherry picking and further filtered for PAINS compounds and known promiscuous compounds in this modality down to 359 compounds (40% inhibition and F-ratio = 0.2-2.0). From there 168 hits were selected for dose-response in the primary TR-FRET assay and after review of the hit structures, a subset (49) of these were prioritized for dry powder purchasing. Of these, 9 compounds reconfirmed as hits, defined as reproducible, converged dose-response fits (IC_50_ <100μM, positive Hill slope) in both retests with at least a 1.5-fold increase in IC_50_ over a counter-screen artifact assay (**Supplementary Table S1**).

Of our prioritized hits we chose to focus on three molecules (SBI-9532, SBI-9635, SBI-2771) that showed potent inhibitory activity in confirmation TR-FRET assays. SBI-9532 is an isatin derivative (**Figure 5A**) with an IC_50_ of 6.2μM (geometric mean of 3.8μM and 10μM, n=2) in the LRP1-BD4-L57 TR-FRET assay (**Figure 5B**). SBI-9635 is also an isatin derivative that varies from SBI-9532 by the N-substitution and the halogen on C5 (**Figure 5C**). SBI-9635 competed in the LRP1-BD4-L57 TR-FRET assay with an IC_50_ of 11μM (geometric mean of 7μM and 18μM, n=2) (**Figure 5D**). Finally, SBI-2771 is a thiazolidinone urea (**Figure 5E**) that competed with an IC_50_ of 4.5μM (geometric mean of 3.2μM and 6.4μM, n=2) (**Figure 5F**). To rule out artifactual activity arising from compound fluorescence, all three were counter-screened for intrinsic fluorescence under assay conditions and showed no significant fluorescent signal (**Supplementary Figure 5A**), supporting the conclusion that their inhibitory activity reflects genuine disruption of the LRP1-BD4 interaction rather than optical interference.

**Figure 5.**
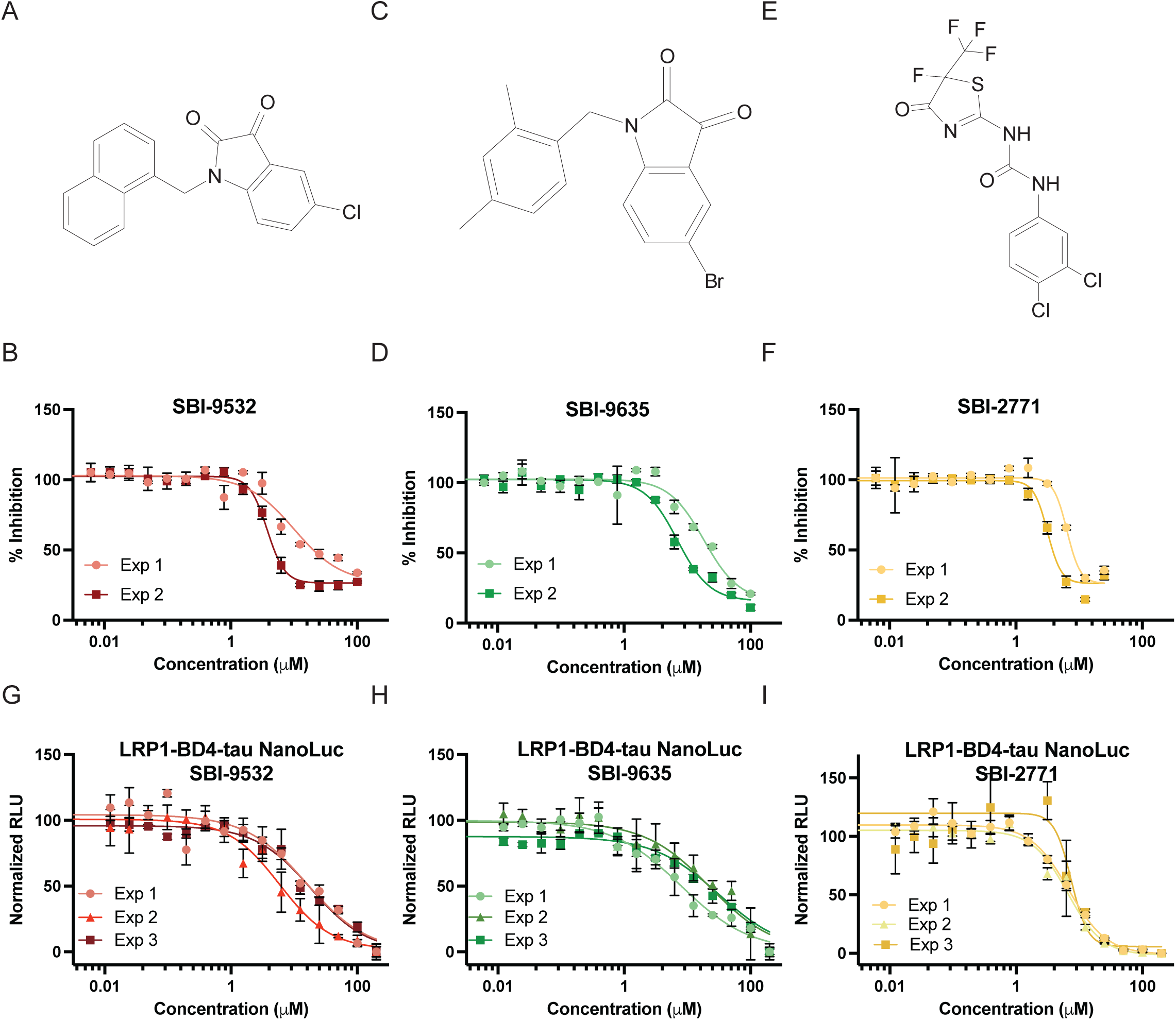
TR-FRET assay validates small molecule inhibitors of the LRP1-BD4-ligand interaction. (A) Chemical structure of SBI-9532 (B) Dose-response curve for two independent replicates of SBI-9532 in the LRP1-BD4-L57 TR-FRET assay; (mean±SD, technical duplicates). (C) Chemical structure of SBI-9635 (D) Dose-response curve for two independent replicates of SBI-9635 in the LRP1-BD4-L57 TR-FRET assay; (mean±SD, technical duplicates). (E) Chemical structure of SBI-2771 (F) Dose-response curve for two independent replicates of SBI- 2771 in the LRP1-BD4-L57 TR-FRET assay; (mean±SD, technical duplicates). (G) Dose-response curve for three independent replicates of SBI-9532 in the LRP1-BD4-tau NanoLuc assay; (mean±SD, technical triplicates). (H) Dose-response curve for three independent replicates of SBI-9635 in the LRP1-BD4-tau NanoLuc assay; (mean±SD, technical triplicates). (I) Dose-response curve for three independent replicates of SBI-2771 in the LRP1-BD4-tau NanoLuc assay; (mean±SD, technical triplicates).

The three priority compounds were subsequently profiled using our LRP1-BD4-tau NanoLuc assay and LRP1-BD4-L57 FP assay to orthogonally confirm activity. All three compounds showed dose-dependent competition of the LRP1-BD4-tau interaction in the NanoLuc assay with IC_50_ values of 12μM (95% CI: 3-56μM, n=3; **Figure 5G**) for SBI-9532, 18μM (95% CI: 3- 105μM, n=3; **Figure 5H**) for SBI-9635, and 6.9μM (95% CI: 5.8-8.2μM, n=3; **Figure 5I**) for SBI- 2771. Surprisingly, none of the three compounds showed measurable competition in the LRP1- BD4-L57 FP assay within the concentration range tested (**Supplementary Figure 5B**). This likely reflects the higher LRP1-BD4 concentration required for this assay format relative to the TR-FRET platform (100nM vs. 30nM) and further emphasizes the utility of having multiple orthogonal assays (**see Discussion**). Together, these three compounds represent a set of validated small molecule inhibitors of the LRP1-BD4-ligand interaction that could then be tested in cellular assays to determine their ability to disrupt tau endocytosis.

### Small-molecule inhibitors of the LRP1-BD4-tau interaction reduce tau uptake in cells

To determine whether compounds identified in biochemical assays could modulate tau biology in a cellular context, we evaluated compound efficacy using a previously established tau uptake assay^6,26^. This assay is performed by taking fluorescently labeled tau (tau-647) and measuring cellular uptake by placing known concentrations into the media of H4 neuroglioma cells. After a one-hour incubation at 37°C, cells are trypsinized and uptake is detected using flow cytometry. This assay is dose-dependent, with monomeric tau-647 displaying an EC_50_ of 41nM (95% CI: 29-58nM, n=3) (**Figure 6A**). As an initial validation, we confirmed that both unlabeled tau and the L57 peptide could competitively inhibit uptake of tau-647 (25nM) when added concurrently to the media (**Figure 6B**), consistent with shared binding to LRP1. Unlabeled tau competed with an IC_50_ of 120nM (95% CI: 50-290nM, n=3) and L57 competed with an IC_50_ of 140nM (95% CI: 100-200nM, n=3).

**Figure 6.**
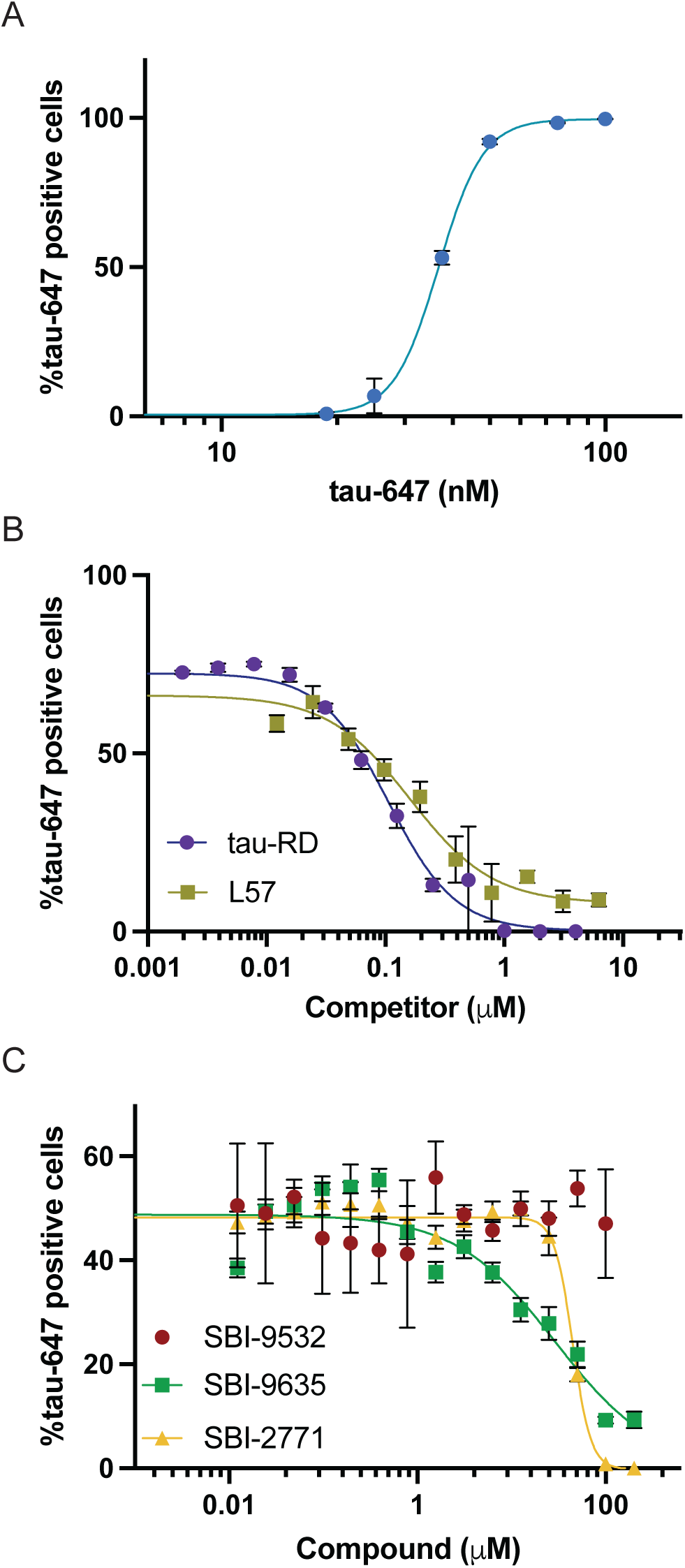
Small-molecule inhibitors of LRP1-BD4 reduce tau uptake in H4 neuroglioma cells. (A) Dose-dependent uptake of tau-647 in H4 cells. Cells were incubated with increasing concentrations of tau-647 for 1h at 37°C and uptake was quantified by flow cytometry, representative experiment is shown (mean±SD, technical triplicates), EC_50_ = 41nM (95% CI: 29- 58nM, n=3 independent experiments) (B) Competitive inhibition of tau-647 (25nM) uptake by unlabeled tau and L57 peptide, representative experiment is shown (mean±SD, technical triplicates). Unlabeled tau IC_50_ = 120nM (95% CI: 50-290nM, n=3 independent experiments) and L57 IC_50_ = 140nM (95% CI: 100-200nM, n=3 independent experiments). (C) Inhibition of tau-647 (25nM) uptake by SBI-9532, SBI-9635, and SBI-2771, representative experiment is shown (mean±SD, technical triplicates). SBI-9635 IC_50_ =19μM (95% CI: 8-46μM, n=3 independent experiments) and SBI-2771 IC_50_ = 54μM (95% CI: 36-81μM, n=3 independent experiments).

Having established this benchmark, we next assessed whether our three lead compounds (SBI- 9532, SBI-9635, SBI-2771) could reduce tau endocytosis in this platform. Holding tau-647 at 25nM we titrated each compound down from 100μM. We found that SBI-9532 was a poor inhibitor of tau-647 endocytosis and did not reproducibly show any inhibition, whereas both SBI- 9635 and SBI-2771 displayed reproducible inhibition profiles. SBI-9635 exhibited an IC_50_ of 19μM (95% CI: 8-46μM, n=3) and SBI-2771 displayed an IC_50_ of 54μM (95% CI: 36-81μM, n=3) (**Figure 6C**). No cellular toxicity was observed for any of the three compounds (**Supplementary Figure 6**). These findings indicate that disruption of the LRP1-BD4-tau interaction *in vitro* translates to functional inhibition of tau uptake in cells. Further, these compounds provide starting points for additional medicinal chemistry efforts to enhance drug-like properties.

### Conclusion

The propagation of tau pathology through interconnected brain regions is a central driver of neurodegeneration in Alzheimer’s disease^1,3,5^, yet the molecular mechanisms underlying tau’s intercellular transfer remain largely untargeted by current therapeutic strategies. LRP1 has emerged as a key mediator of tau uptake^6,7^, but the biochemical targeting of this interaction has been constrained by the challenges of recombinant LRP1 production and the absence of quantitative, screening-compatible binding assays with LRP1 ligands. In this study, we address these limitations through the recombinant production of LRP1-BD4, the development of three orthogonal ligand binding assay platforms, and the execution of high-throughput screening campaigns that identify small-molecule inhibitors capable of reducing tau uptake in cells.

We focused our initial efforts on developing methodology to express and purify the fourth ligand binding domain fragment of LRP1 (LRP1-BD4), a region of LRP1 that we previously had shown to be necessary for tau endocytosis^6^ and which has been implicated in binding to other known LRP1 ligands^27^. The successful recombinant expression and purification of a functional LRP1- BD4 protein represents an enabling technical advance. The BD4 domain of LRP1 is composed of 11 complement-type repeats stabilized by 33 intradomain disulfide bonds and extensive calcium coordination throughout^19,28^, making it a formidable challenge for expression. By co-expressing LRP1-BD4 with its endogenous chaperone RAP^29^, we were able to produce sufficient quantities for both our biochemical studies and two high-throughput screening efforts. We used biophysical characterization to analyze our purified LRP1-BD4 protein and found that it had a melting temperature of 68°C and a CD spectrum matching structural predictions. The robustness of our expression system now makes LRP1-BD4 accessible for broader biochemical and structural studies, including potential cryo-EM efforts with bound ligands or inhibitors.

A central finding of this work is the development and application of three orthogonal assay formats to characterize LRP1-ligand interactions. Using fluorescence polarization, split luciferase complementation, and TR-FRET, we consistently demonstrated that tau, L57, and RAP are competitive binding partners for LRP1-BD4. Our cross-format validation is significant because each assay format is susceptible to distinct artifact profiles: FP can be affected by compound interference^30,31^; luminescence-based assays can be influenced by compound effects on enzyme activity or protein stability^32–35^; and TR-FRET, while more resistant to optical interference than traditional FRET, can still be perturbed by compounds that affect donor-acceptor distance independent of binding site competition^36,37^. The convergence of competition data across these platforms strengthens the conclusion that tau, L57, and RAP engage overlapping or mutually exclusive binding surfaces on BD4.

However, several nuances warrant consideration. Our split luciferase assay requires productive LgBiT-SmBiT complementation in addition to binding, which would be expected to yield an apparent affinity weaker than the true K_d_, as not all binding events may produce the correct tag orientation for luminescence. The developed FP assay measures L57 binding rather than tau binding directly, and the relationship between the L57 and tau binding epitopes on BD4, while shown to be competitive, may not be fully overlapping. The crystal structure of RAP bound to LDLR suggests that binding to LRP1 is likely mediated by interaction across multiple CRs^38^. Therefore, RAP competes with both L57 and tau, but could do so through steric occlusion rather than direct site overlap and indeed we did observe that RAP was not capable of fully competing our L57 tracer in the FP platform. Resolving the precise binding epitopes through structural studies would clarify these relationships and inform rational inhibitor design.

The two TR-FRET screening campaigns illustrate a fundamental challenge in screening against protein-protein interactions involving intrinsically disordered proteins (IDPs). The tau-based TR-FRET screen, which directly measured the LRP1 ligand interaction of interest, yielded a high hit rate (2.2%) with strong re-confirmation. Rather than reflecting stochastic false positives, this pattern is consistent with systematic artifacts arising from tau’s intrinsic disorder and compounds that promote tau aggregation or non-specifically displace tau from BD4 would be reproducibly active yet mechanistically irrelevant. This is a recognized challenge in screening against IDP-containing interactions, where the conformational plasticity of the disordered partner creates multiple compound-sensitive nodes that do not correspond to the physiological binding interface^39,40^. We therefore developed the L57-based TR-FRET screen as our primary screening format, reasoning that the small peptide tracer would be less susceptible to these artifacts while still reporting on the same competitive binding interface on BD4. The lower hit rate (0.98%) and the subsequent identification of compounds with cellular activity against tau endocytosis support this rationale. Further consistent with this notion, both of our TR-FRET screens sampled the same 10,240 compound Chembridge library and upon cross-referencing we found a low overlap of hits; of 232 hits in LRP1-BD4-tau only 20 were hits in LRP1-BD4-L57 (8.6%). We interpret this as a substantial fraction of our tau hits representing potential artifacts due to tau’s conformational disorder rather than genuine competitive inhibitors of the BD4 interface. However, this approach carries a trade-off: the L57 format reports on disruption of the L57-BD4 interface as a proxy for the tau-BD4 interface, and compounds targeting tau-specific binding determinants not shared with L57 would be missed. Future campaigns could benefit from a tiered strategy using the L57 format as a primary screen followed by secondary profiling in the tau format to capture both classes of inhibitors.

An additional consideration is that apparent competitive potency of inhibitors may depend on the protein concentrations used in a given assay format. The L57 FP assay requires a higher LRP1- BD4 concentration (100nM LRP1-BD4) than the L57 HTRF (30nM LRP1-BD4) or tau NanoLuc formats (20nM LRP1-BD4), and in the L57 FP assay, the LRP1-BD4 concentration exceeds the measured K_d_ of 42nM. Under these conditions, weaker competitors would be expected to show a right-shifted apparent IC_50_, which could explain why our priority compounds, which were active in the L57 HTRF and tau NanoLuc assays, did not show measurable competition in the FP format. This observation underscores the value of employing multiple, complementary assay formats when evaluating primary hits, since apparent activity can be shaped by assay-specific parameters independent of a compound’s true target engagement.

The cellular activity of SBI-9635 and SBI-2771, while modest (IC_50_ values of 19μM and 54μM, respectively), demonstrates that biochemical inhibition of the LRP1-BD4-ligand interaction can translate to functional reduction of tau endocytosis. The attenuation of potency from biochemical to cellular assays is consistent with additional barriers present in a cellular context, including serum proteins that can reduce free-compound^41^ or the dynamic endocytic cycling of LRP1 that could limit the available receptor pool^42^. Notably, SBI-9532, which showed potent biochemical activity (IC_50_ = 6.2μM), failed to reproducibly inhibit cellular tau uptake, highlighting the well-recognized disconnect between biochemical and cellular activity and underscoring the importance of cellular validation early in hit-to-lead campaigns.

Several important limitations should be acknowledged. First, our cellular assays were performed in H4 neuroglioma cells, which, while a well-established model for tau uptake studies, do not fully recapitulate the neuronal context in which tau propagation occurs *in vivo*. Future studies should evaluate compound efficacy in primary neurons^43^ and in *in vivo* models of tau spread^6,44^. Second, determining whether our compounds act on-target through LRP1 in cells presents a methodological challenge: LRP1 knockdown eliminates tau uptake entirely, precluding the standard genetic validation approach of testing compound efficacy in the absence of the target. The lack of endocytosis in the absence of LRP1 confirms that LRP1 is the dominant mediator of tau internalization under our assay conditions, but it leaves open the possibility that compounds could reduce uptake through LRP1-independent mechanisms, such as interference with downstream endocytic machinery or non-specific membrane effects. Several approaches could address this question, including partial LRP1 knockdown to shift the dynamic range of uptake or correlation of biochemical and cellular potency across an expanded compound series. These experiments are priorities for ongoing validation. Third, the selectivity of our compounds against other LRP1 ligand interactions remains uncharacterized. Given LRP1’s broad ligand repertoire^9^ and its roles in lipid metabolism^45^, protease clearance^46^, and signaling^47^, any therapeutic strategy targeting this receptor must be carefully assessed for potential off-pathway effects. Lastly, the mechanism by which our compounds disrupt the BD4-ligand interaction, whether through direct competition at the binding site, allosteric modulation, or disruption of calcium coordination, remains to be determined.

From a broader perspective, this work contributes to an emerging recognition that endocytic receptors represent tractable targets for modulating pathological protein uptake in neurodegenerative diseases. Proteins such as α-synuclein have been reported to spread in disease and several receptors have been proposed^48–50^, suggesting that the assay development and screening framework established here could be generalized to other protein-receptor pairs. Moreover, our experience with the tau-based HTS highlights the particular challenges of screening against IDP-containing interactions and suggests that structured surrogate tracers, while imperfect, may offer a more tractable path to identifying genuine inhibitors.

In summary, we have established a comprehensive biochemical and screening platform for targeting the tau-LRP1 interaction and identified small-molecule inhibitors that reduce tau uptake in cells. While the current compounds lack the potency and selectivity profiles needed for therapeutic leads, they provide validated starting points for medicinal chemistry optimization. More broadly, this work demonstrates that the tau-LRP1 interface is pharmacologically accessible, and it provides the tools and framework necessary to pursue this target with increasing rigor and sophistication.

## Materials and Methods

### Protein expression and Purification

Tau constructs used in this study include the longest CNS isoform of tau (2N4R), 2N4R fused to smBiT tag (2N4R-smBiT and smBiT-2N4R) and a truncated tau construct spanning the microtubule-binding repeat domain (tau-RD; residues 244-372). Tau constructs were expressed in One Shot™ BL21 Star™ (DE3) cells (Invitrogen) and purified as previously described^26,51^. Tau protein was labeled with AlexaFluor-647 as previously described^6,26^. Average degree of labeling was ∼1.

RAP constructs were expressed in One Shot™ BL21 Star™ (DE3) cells (Invitrogen) and purified as previously described^6,51^.

LRP1-BD4 was expressed in Expi293F™ cells (Thermo Fisher Scientific) with a co-transfection of a mammalian construct of RAP, according to the manufacturer’s instructions (Gibco). Cells were maintained at 37°C with 8% CO_2_ for 5 days, after which cultures were harvested by centrifugation at 1,000xg for 10 min. The clarified supernatant was collected for purification. Ni-NTA resin (Thermo Fisher Scientific) was equilibrated by washing twice with Milli-Q water followed by two washes with His binding buffer (50mM HEPES, pH 8.0, 1M NaCl, 10mM imidazole), centrifuging at 2,000xg for 3 min between washes. The clarified supernatant was incubated with the equilibrated resin, and the total volume was adjusted to 40mL with His binding buffer. The mixture was rotated overnight at 4 °C. Resin was loaded onto a gravity-flow column and washed with 10mL of His binding buffer, followed by three washes with 10mL of His wash buffer (50mM HEPES, pH 8.0, 300mM NaCl, 30mM imidazole). Bound protein was eluted by incubating the resin with 5mL of His elution buffer (50mM HEPES, pH 8.0, 300mM NaCl, 300mM imidazole) for 15 min prior to collection. An additional 5 mL elution was collected to maximize recovery. Eluted protein was dialyzed overnight against high-EDTA buffer (50 mM HEPES, pH 8.0, 20 mM NaCl, 100 mM EDTA) prior to further purification by anion exchange chromatography using an ÄKTA Pure system (Cytiva). The column was equilibrated with water followed by Buffer A (50mM HEPES, pH 8.0, 20mM NaCl, 20mM EDTA). The sample was applied at 0.5 mL/min, and nonspecifically bound proteins were removed by washing with 3–5 column volumes of Buffer A at 5 mL/min. Protein was eluted using a linear gradient from 0– 100% Buffer B (50mM HEPES, pH 8.0, 1M NaCl, 20mM EDTA) over 124 mL. Fractions were collected and analyzed by SDS-PAGE, and those containing pure protein were pooled. The purified protein was concentrated using 10 kDa molecular weight cutoff Amicon filters and buffer-exchanged into storage buffer (20mM HEPES, pH 8.0, 150mM NaCl, 1mM CaCl_2_). Protein concentration was determined via nanodrop, aliquoted, flash-frozen in liquid nitrogen, and stored at −80 °C.

For removal of the MBP tag, purified protein was exchanged into thrombin cleavage buffer (20mM Tris-HCl, pH 8.4, 150mM NaCl, 2.5mM CaCl_2_) and incubated with thrombin (30 U per mg protein) overnight at 4 °C with gentle agitation. Following cleavage, the sample was buffer-exchanged into maltose binding buffer (20 mM HEPES, pH 8.0, 20 mM NaCl) and applied to an MBPTrap™ HP column. The cleaved BD4 protein was collected in the flowthrough during sample application and validated by SDS-PAGE.

### Circular Dichroism

Thrombin cleaved LRP1-BD4 sample (4uM) in a 5mM phosphate buffer was measured on a Jasco J-1500 Circular Dichroism Spectrophotometer. Readings were taken from 200nm to 260nm at room temperature. Data was analyzed using Jasco’s Secondary Structure Estimation Calibration Model and Circular Dichroism Multivariate Secondary Structure Estimation Program to determine relative secondary structure.

### Differential Scanning Fluorimetry (DSF)

DSF experiments were performed in clear, low-volume 384-well qPCR plates (Thermo Fisher Scientific) with a total volume of 20µL per well. Each reaction contained 2.5 µM MBP-cleaved LRP1-BD4 and 50µM MWF08 dye^52^ and was analyzed using a QuantStudio 5 Real-Time PCR System (Thermo Fisher Scientific). Samples were heated from 25°C to 99°C at a rate of 0.05°C/s, with fluorescence measurements collected in the FAM channel (Ex: 470 nm, Em: 520 nm). Fluorescence data were exported and analyzed in GraphPad Prism. Data were fit to a four-parameter logistic model to determine melting temperatures (Tm), defined as the midpoint of the transition.

### Fluorescence Polarization Assay

Purified LRP1-BD4 protein was thawed on ice and diluted in FP assay buffer (20mM HEPES, pH 8.0, 150mM NaCl, 1mM CaCl_2_, 0.05% CHAPS). Assays were performed in black, low-volume 384-well plates (Corning) utilizing 20nM final FITC-labeled L57 peptide (Genscript). Control wells containing LRP1-BD4, but lacking probe were included for background subtraction. Plates were incubated at room temperature for 30 min, and read on a SpectraMax i3x plate reader (Molecular Devices) with excitation at 485 nm and emission at 535 nm. Parallel (p) and perpendicular (s) fluorescence intensities were recorded, and background-subtracted values were used to calculate fluorescence polarization (FP). FP signal at 0µM LRP1-BD4 was subtracted to determine the ΔmP values. Data were analyzed in GraphPad Prism (v11), and binding curves were fit by nonlinear regression using a four-parameter logistic model to determine apparent K_d_ values.

For competition experiments, competitors were serially diluted and combined with LRP1-BD4 (0.1µM or 1µM) and FITC-L57 tracer (20nM) in 384-well plates as described above. Control wells containing competitor dilutions, but lacking probe were included for background subtraction. Control experiments lacking LRP1-BD4 but including tracer and competitor were also run to rule-out non-specific tracer binding. After incubation and measurement, FP values were calculated and analyzed as above, with FP signal at 0µM LRP1-BD4 subtracted to determine the ΔmP values. Data were analyzed in GraphPad Prism (v11), and binding curves were fit by nonlinear regression using a four-parameter logistic model to determine IC_50_ values.

### Split Complementation Assay

Split luciferase complementation assays were performed in white, half-area 96-well plates (Greiner Bio-One) in phosphate-buffered saline (PBS; Gibco) supplemented with 0.01% Triton X-100. Dilutions of tau-SmBiT were incubated with MBP-LgBiT-LRP1-BD4(10nM) or MBP-LgBiT (10nM) at room temperature for 10 min to allow complex formation. Furimazine (Lumiprobe) was then added to a final concentration of 10μM, followed by an additional 10 min incubation at room temperature. Luminescence was measured using a SpectraMax i3x plate reader (Molecular Devices). Data were analyzed in GraphPad Prism (v11), and binding curves were fit by nonlinear regression using a four-parameter logistic model to determine EC_50_ values.

Competition assays were performed to evaluate the ability of unlabeled ligands to disrupt the LRP1-BD4-tau interaction. Reactions contained 20nM MBP-LgBiT-LRP1-BD4, 0.2µM SmBiT- tau, and serial dilutions of competitors. Control wells lacking SmBiT-tau were included to define baseline signal. MBP-LgBiT-LRP1-BD4 was first combined with competitors, followed by addition of tau-SmBiT. Plates were incubated at room temperature for 10 min. Furimazine (Lumiprobe) was added (10μM), and plates were incubated for an additional 10 min prior to luminescence measurement on a Spectramax i3x plate reader. Data were analyzed in GraphPad Prism (v11), and binding curves were fit by nonlinear regression using a four-parameter logistic model to determine IC_50_ values

## TR-FRET Assay

### LRP1-BD4-tau TR-FRET assay

LRP1-BD4-tau time resolved Förster resonance energy transfer (TR-FRET) assays were performed in black, flat-bottom 1536-well format. An Eu^3+^-labeled anti-Flag antibody (Revvity) served as the fluorescence donor, recognizing Flag-tagged LRP1-BD4, while tau labeled with Alexa Fluor-647 (tau-647) served as the acceptor. Control wells containing only one protein partner were included as positive low-signal controls. Plates were incubated at room temperature for 1hr before and after additional of the donor, and read on a PHERAstar plate reader (BMG Labtech) with excitation at 337nm and emissions at 620nm and 665nm. An Eu^3+^- labeled anti-MBP (Revvity) donor was also tested for this assay but did not yield detectable binding signal, potentially due to the donor-acceptor pair falling outside the effective FRET range for this configuration.

### LRP1-BD4-L57 TR-FRET assay

LRP1-BD4-L57 TR-FRET assays were performed in black, flat-bottom 384-well format. An Eu^3+^- labeled anti-MBP antibody (Revvity) served as the fluorescence donor, recognizing MBP-tagged LRP1-BD4, while Biotin-L57 recognized by streptavidin-D2 served as the acceptor. Control wells containing only one partner were included as positive low-signal controls. Plates were incubated at room temperature for 1 hr before and after addition of the donor-acceptor mix, and read on a PHERAstar plate reader (BMG Labtech) with excitation at 337nm and emissions at 620nm and 665nm. An Eu^3+^-labeled anti-FLAG (Revvity) donor was also tested for this assay but did not yield detectable binding signal, potentially due to the donor-acceptor pair falling outside the effective FRET range for this configuration.

### High-Throughput Screening Pilot screen LRP1-BD4-tau

The LRP1-BD4-tau TR-FRET assay was used to screen a library of 10,240 small molecules from the Chembridge Library collection in 384-well format. Compounds were tested at a single concentration of 25 µM. tau-647 and LRP1-BD4 were held constant at 5nM. Each plate included positive control wells (no LRP1-BD4) and negative control wells (DMSO vehicle).

### Primary screen LRP1-BD4-L57

The LRP1-BD4-L57 TR-FRET assay was used to screen a library of 150,000 small molecules (100,000 Enamine Discovery Diversity library and 50,000 Chembridge Library collection) in 1536-well format. Compounds were tested at a single concentration of 25 µM, with LRP1-BD4 and Biotin-L57 held constant at 30 nM and 200 nM, respectively. Each plate included positive control wells (no Biotin-L57) and negative control wells (DMSO vehicle). The screen exhibited excellent performance metrics (signal-to-background ratio S/B = 5.68 ± 0.26; Z-factor = 0.81 ± 0.03). An initial hit rate of 0.98% (1,476 compounds) was achieved using a primary threshold of ≥40% inhibition. Primary hits were confirmed by cherry-pick retesting in duplicate or triplicate in the same TR-FRET assay format; confirmed hits were defined as those achieving ≥40% inhibition in replicate testing.

### Hit triage and dose-response

Confirmed hits were subjected to manual medicinal chemistry review to remove pan-assay interference compounds (PAINS), reactive functional groups, and known promiscuous inhibitors in fluorescence-based assays. An F-ratio filter (0.2–2.0) was additionally applied to exclude compounds with intrinsic fluorescence that could artifactually suppress TR-FRET signal. This triage reduced the confirmed hit set to 359 compounds. From these, 168 compounds were selected for dose-response analysis in the primary LRP1-BD4-L57 TR-FRET assay. Compounds were tested across a 5-point, 2-fold serial dilution series from a top concentration of 100 µM. IC_50_ values were calculated by fitting normalized inhibition data to a four-parameter logistic (4PL) model. Based on dose-response IC_50_ values and further medicinal chemistry assessment of compound structures, 49 compounds were prioritized for dry powder purchase and follow-up characterization. Powder compounds were re-tested in duplicate dose-response in the primary LRP1-BD4-L57 TR-FRET assay, alongside a counter-screen artifact assay (biotinylated MBP-TEV-Thrombin-FLAG-HA-Avi-His construct using Eu^3+^-anti-MBP and streptavin-D2). A compound was considered a confirmed hit if both replicate dose-response curves converged to a fit with a positive Hill slope (indicating inhibition), an IC_50_ < 100µM, and if the artifact assay showed at least a 1.5-fold higher IC_50_ than the mean primary-assay IC_50_. Nine compounds met these criteria, and of these three compounds were prioritized for further testing based on the largest fold-separation between primary and artifact assays. Compounds were purchased from Molport and purity was assessed via LC/MS (**Supplementary Figure 7**).

### Cellular Tau Uptake

H4 neuroglioma cells were cultured in DMEM-10 media (DMEM + 10% FBS, 1% Pen/Strep) and seeded in tissue culture-treated 96-well plates (Corning) at a density of 5 × 10^4^ cells per well and incubated overnight at 37 °C in a humidified atmosphere containing 5% CO_2_.On the day of the assay, culture medium was removed and cells were washed once with phosphate-buffered saline (PBS; Gibco). Prewarmed DMEM-10 (45 μL) was added to each well, followed by addition of Alexa Fluor 647–labeled tau (tau-647; 5 μL per well of the appropriate working solutions). For tau titration experiments, tau-647 was added at increasing concentrations as indicated. Plates were gently mixed and incubated for 1 h at 37 °C in a humidified atmosphere containing 5% CO_2_. Following incubation, the uptake solution was aspirated, and cells were detached using 100μL of 25% trypsin in PBS for 4 min at 37 °C. Trypsinization was quenched by addition of 100 μL of 25% fetal bovine serum (FBS) in PBS containing propidium iodide for live/dead discrimination. Cells were analyzed by flow cytometry, and propidium iodide positive cells were excluded from analysis. Tau uptake was quantified based on Alexa Fluor 647 fluorescence intensity in viable cells. Cells incubated in the absence of tau-647 served as negative controls for gating and background fluorescence determination.

For compound competition experiments, cells were treated with a fixed concentration of tau-647 in the presence of test compounds added concurrently at the indicated concentrations. Plates were gently mixed and incubated for 1 h before processing as described above.

### Supporting Information

SDS-PAGE and circular dichroism characterization of purified LRP1-BD4; L57 peptide self-competition, L57A negative control, and RAP-L57-FITC binding control for the fluorescence polarization assay; SmBiT-tag orientation and competition controls for the NanoLuc complementation assay; TR-FRET assay specificity controls and cherry-pick reconfirmation of tau-based screening hits; intrinsic fluorescence counter-screen, fluorescence polarization, cellular toxicity and LC/MS purity data for the three priority compounds; table of the dose-response and counter-screen artifact-assay data for prioritized compounds.

## Acknowledgements

This work was supported by grants from the National Institutes of Health (R01 AG075084 to K.S.K. & M.R.J., R01 AG077672 to J.N.R.), Paul Slavik (J.N.R.), the Alzheimer’s Drug Discovery Foundation (K.S.K.), the University of Massachusetts IALS Translational Fellowship (C.W.). We thank Dr. Jason Gestwicki (UCSF) for the MWF08 dye used in DSF experiments. We thank Ian Pass for his guidance with the HTS design. We are thankful to Dr. Steve Eyles and the University of Massachusetts Amherst Core Facilities for use of the Mass Spectrometry Core (RRID: SCR_019063).

