## Supplemental Information for "High-Throughput Screening Identifies Small-Molecule Inhibitors of the Tau-LRP1 Interaction"

**Supporting Figure Legends**

**Figure S1.** (A) SDS-PAGE Coomassie gel of MBP-LRP1-BD4 before (Lane 1) or after cleavage by thrombin (Lane 2 – LRP1-BD4; Lane 3 – MBP). (B) Alphafold3 prediction of LRP1-BD4. Model is colored by per-residue confidence score (pLDDT): dark blue (>90, very high), cyan (70-90, confident), yellow (50-70, low), and orange (<50, very low). Inset: Predicted Aligned Error (PAE) plot, where darker green indicates lower expected positional error (Å) between residue pairs.

**Figure S2.** (A) Competition of unlabeled L57 with FITC-L57 for LRP1-BD4 binding. LRP1-BD4 (100nM) with L57-FITC (20nM), three independent experiments shown (mean±SD, technical triplicates) IC_50_ = 156nM (95% CI: 104-234nM, n=3 independent experiments). (B) Competition of unlabeled L57A with FITC-L57 (20nM) for LRP1-BD4 (100nM) binding, three independent experiments shown (mean±SD, technical triplicates). (C) RAP titration with L57-FITC (20nM), two independent experiments shown (mean±SD, technical triplicates).

**Figure S3.** (A) Titration of tau-SmBiT (C-term) or SmBiT-tau (N-term) with 10nM MBP-LgBiT-LRP1-BD4 produces luminescent signal (mean±SD, technical triplicates). Three representative experiments shown. (B) Representative titration of N-term SmBiT-tau (20nM) with tau-RD, L57, or RAP indicates competitive binding (two independent experiments, mean±SD, technical triplicates).

**Figure S4.** (A) LRP1-BD4-tau-647 TR-FRET competition with unlabeled tau and L57. Tau IC_50_ = 3.3nM (95% CI: 2.9-3.9nM) and L57 IC_50_ = 36nM (95% CI: 23-66nM) (B) Hits from LRP1-BD4-tau TR-FRET screen show similar % Activity in the HTS and in the reconfirmation assay. (C) LRP1-BD4-L57 TR-FRET competition with unlabeled L57, tau, and L57A. L57 IC_50_ = 106nM (95% CI: 71-167nM), unlabeled tau IC_50_ = 66nM (95% CI: 50-87nM), and L57A was non-competitive.

**Figure S5.** (A) SBI-9532, SBI-9635, and SBI-2771 show minimal activity in the artifact assay. (B) SBI-9532, SBI-9635, and SBI-2771 do not compete in the L57 FP assay.

**Figure S6.** Viability of H4 neuroglioma cells with increasing concentrations of SBI-9532, SBI-9635, and SBI-2711.

**Figure S7.** LC/MS of SBI-9532, SBI-9635, and SBI-2771.

**Table S1.** Dose-response and artifact assay of purchased compounds.

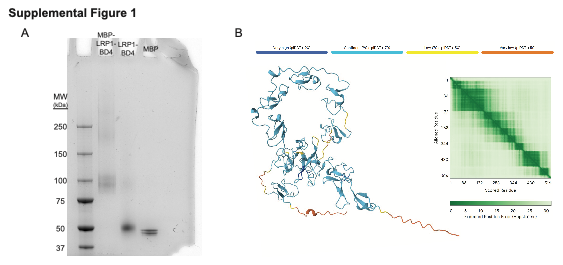

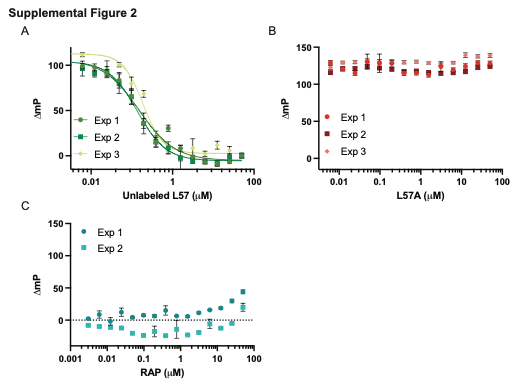

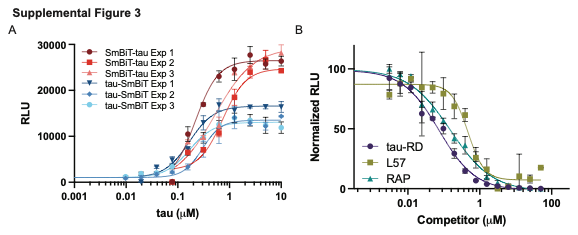

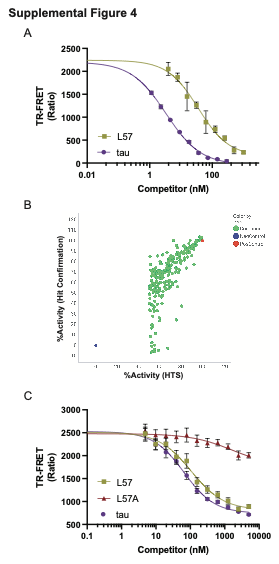

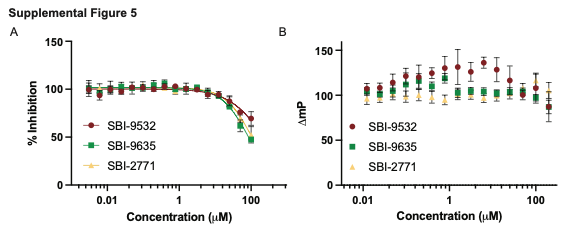

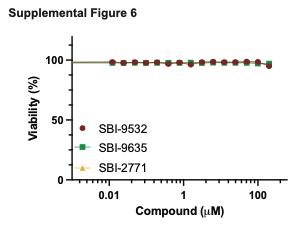

**Supplemental Figure 7**

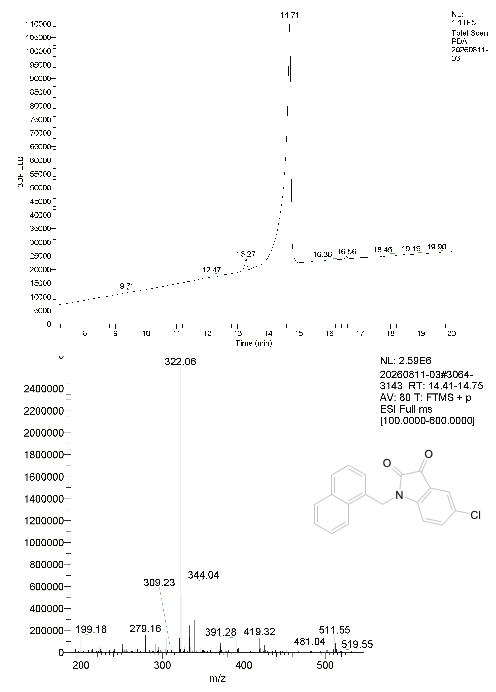

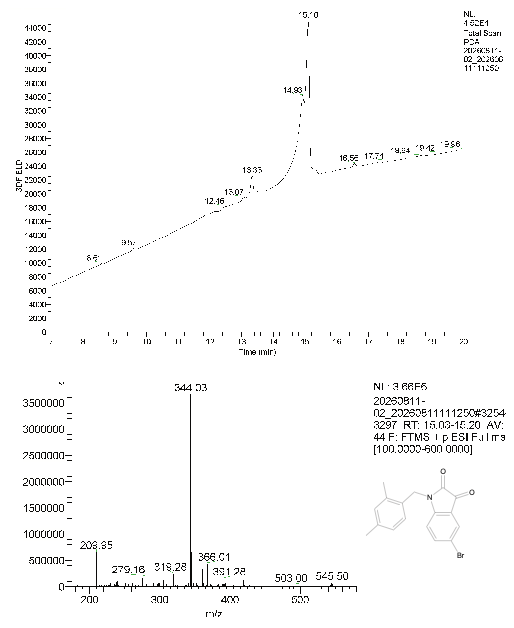

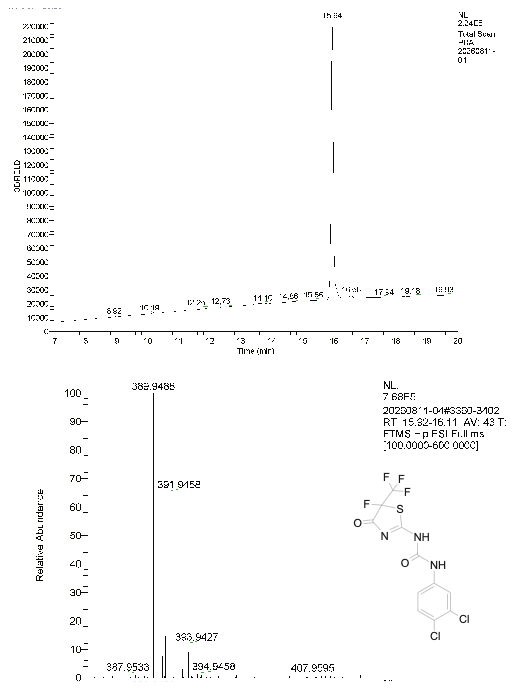
